# Enhancing Clinical Classification of Protein Variants using ESM2 and UMAP

**DOI:** 10.1101/2025.07.26.666924

**Authors:** Ugo Lomoio, Pierangelo Veltri, Pietro Hiram Guzzi

## Abstract

Protein sequences may vary due to mutations in their coding DNA sequence, leading to differences in structure and function. The same protein may exist in multiple variant forms, each potentially leading to distinct phenotypic consequences depending on how the alterations affect its structure, function, or expression. Missense variants are single nucleotide substitutions in the DNA sequence that result in the replacement of one amino acid with another in the corresponding protein, potentially altering its structure, stability, or function. The clinical interpretation of missense variants in protein-coding regions remains a fundamental challenge in genomic medicine. Recent advances in protein language models and manifold learning provide new opportunities for unsupervised extraction of biologically relevant information from protein sequences. In this work, we integrate representations derived from ESM2 (**spiegare**) with nonlinear dimensionality reduction via UMAP (**spiegare**) to improve the classification of variants of uncertain significance (VUS) in disease-associated proteins. Our results suggest that this approach improves separability of benign and pathogenic variants, offering a scalable and interpretable strategy for variant prioritization in precision medicine.

## 1 Introduction

The growing accessibility of genome sequencing technologies has ushered in a new era of precision medicine, enabling the identification of genetic variants at an unprecedented scale. However, while sequencing pipelines can detect thousands of variants per individual, interpreting their clinical relevance remains a formidable challenge. Among protein-coding alterations, missense variants — those resulting in amino acid substitutions — are particularly difficult to interpret due to the complex relationship between sequence variation, protein structure and function, and clinical phenotype [5]. As a result, a large proportion of such variants are classified as variants of uncertain significance (VUS), which poses significant limitations to genetic diagnosis, risk assessment, and personalized treatment [8].

Traditional computational tools for variant effect prediction, such as SIFT, PolyPhen-2, CADD, and REVEL, leverage features based on evolutionary conservation, sequence homology, protein structure, and functional annotations [1]. While these tools provide valuable insights and have been widely adopted in clinical practice, they are constrained by limitations in interpretability, gene-specific biases, and generalizability across variant classes. Moreover, their reliance on curated databases or explicit alignments restricts their scalability to rare or novel variants, which often lack sufficient homologous information.

Recent advances in self-supervised learning, particularly in the domain of protein language modeling, have opened new frontiers in computational biology. Inspired by developments in natural language processing, protein language models (PLMs) are trained on vast corpora of unaligned amino acid sequences using transformer-based architectures to learn contextual embeddings that implicitly capture structural, functional, and evolutionary signals [6]. These models, such as ESM (Evolutionary Scale Modeling), ProtBERT, and TAPE, are capable of producing high-dimensional vector representations of protein sequences or individual residues without relying on explicit multiple sequence alignments.

Among these, ESM2 represents the most recent and powerful iteration in the ESM family of models, trained on over 60 billion amino acids from UniRef and comprising up to 15 billion parameters [3]. ESM2 generates contextualized embeddings at the residue and sequence level that have demonstrated state-of-the-art performance across several downstream tasks, including structure prediction, contact map inference, and mutational effect prediction. These embeddings have the potential to encode nuanced information about protein folding, dynamics, and biochemical function, making them highly attractive for variant effect classification.

However, despite the richness of PLM-derived representations, their high dimensionality (e.g., 1,280 dimensions for ESM2-T36) presents significant challenges in terms of visualization, interpretability, and downstream machine learning. Raw embeddings may contain noise, redundancy, or non-linearly separable patterns that complicate classification. To address these limitations, dimensionality reduction techniques are frequently employed to map high-dimensional vectors into lower-dimensional manifolds that preserve essential structural features.

Uniform Manifold Approximation and Projection (UMAP) is a nonlinear dimensionality reduction algorithm that preserves both local and global relationships in data while producing embeddings that are well-suited for visualization and clustering [4]. UMAP has been used successfully in genomics and single-cell transcriptomics, where high-dimensional latent spaces are common. When applied to PLM-derived embeddings, UMAP enables better separation of latent biological classes and enhances the interpretability of variant spaces.

Computational methods have become instrumental in predicting the functional effects of missense mutations, offering valuable insights into personalized medicine [9]. Databases such as ClinVar and DECIPHER, along with matchmakers like GeneMatcher, facilitate the sharing of variants and phenotypic descriptions, aiding in the determination of disease relevance [12]. Assessing genetic intolerance through Homologous Missense Constraint measures can further inform the clinical significance of missense variants associated with human diseases [2]. Evaluating regional intolerance using tools like MTR-Viewer can help identify pathogenic missense variants [7]. These methods highlight functionally significant regions within genes that traditional approaches may overlook [7]. Furthermore, databases incorporating protein contactmaps, sequence-to-structure mapping, and stability predictions enhance variant analysis, while evolutionary fitness effect scoring aids in predicting the deleteriousness of variants, potentially reclassifying variants of unknown significance [13] [11]. However, not all mutations have severe damaging impacts, and most produce subtle effects with unclear clinical significance, necessitating accurate measurement of changes in binding affinity induced by mutations [15]. Functional analyses of protein domains have revealed that specific protein functions are associated with varying sensitivities to mutations, suggesting that future pathogenicity predictors may benefit from considering functional annotation [14]. The identification of critical mutation sites and understanding the complex patterns of mutation effect of viral proteins are also crucial, especially for forecasting dominant variants [10].

## 2 The Proposed Method

### 2.1 Dataset Collection

We constructed a curated dataset of missense variants for multiple amyloidosis related proteins from the ClinVar database (accessed June 2025). Focusing on the following proteins:

- **Alpha-1-antitrypsin (P02768)**, a protease inhibitor that forms toxic polymers in liver cells when mutated, leading to liver disease and predisposing to secondary amyloidosis.
- **Transthyretin (P02766)**, thyroid hormone carrier with over 140 known mutations causing hereditary cardiac and neurological amyloidosis. Also the wild-type can causes agerelated cardiac amyloidosis.
- **Fibrinogen** *α***-chain (P02671)**, a blood clotting protein whose mutations cause renal-limited amyloidosis with rapid progression to kidney failure.
- **Apolipoprotein A-I (P02647)**, High-Density Lipoprotein (HDL) component causing hereditary amyloidosis affecting liver, kidneys, heart, or other organs depending on mutation location.
- **Lysozyme (P61626)**, an antimicrobial enzyme causing rare hereditary amyloidosis with renal, gastrointestinal, and sicca symptoms.
- *β*_2_**-microglobulin (P61769)**, a component of the Major Histocompatibility Complex 1 (MHC-I) that accumulates in dialysis patients, causing joint and bone disease including carpal tunnel syndrome.
- **Cystatin C (P01034)**, a protease inhibitor causing hereditary cerebral amyloid angiopathy with recurrent brain haemorrhages in young patients.

For each protein, we retained the canonical UniProt sequence and mapped variants to their corresponding amino acid positions.

Since we aim to perform a binary classification, variants labelled as “likely benign” or “likely pathogenic” were initially treated and approximated as “benign” and “pathogenic” respectively. Then, with the help of UMAP embeddings, we visualized the position of those “likely” mutations and interpret the prediction.

### 2.2 ESM2 Embedding

We used the pretrained ESM2-T36 model [3], implemented in PyTorch and available through the FAIR/ESM GitHub repository, to compute residue-level embeddings. Each variant sequence was mutated in silico at the position of interest, and the corresponding 1,280-dimensional embedding vector was extracted for the mutated residue. No fine-tuning was performed.

### 2.3 Dimensionality Reduction via UMAP2

To reduce the dimensionality of the ESM2-derived vectors, we applied UMAP2 [4] using the umap-learn package. We selected parameters empirically: `min_dist = 0.0`, `metric = “euclidean”`, and a variable `n_neighbors parameter` to overcome cases of dataset unbalance (i.e. more benign than pathogenic mutation). The reduced embeddings were projected to 2D spaces for visualization and classification. Classification of each variant was performed on their distance from the wild-type: pathogenic if the distance is greater then a given threshold, benign otherwise. We confirmed that pathogenic and benign variants formed separable clusters in the reduced space, and variants of uncertain significance (VUS) typically populated inter-cluster regions.

**Table 1:** Detailed ROC-AUC scores of different classification models for each protein related with amyloidosis.

| Protein | VESM++ | AlphaMissense | ESM2+UMAP | N. Benign | N. Pathogenic |
| --- | --- | --- | --- | --- | --- |
| P02768 | 0.6403 | 0.6747 | **0.6820** | 57 | 15 |
| P02766 | 0.7720 | 0.6978 | **0.8941** | 14 | 126 |
| P02671 | 0.6002 | 0.7063 | **0.8098** | 34 | 28 |
| P02647 | - | - | **0.7762** | 22 | 42 |
| P61626 | **0.8611** | - | 0.6667 | 11 | 6 |
| P61769 | - | - | **0.6667** | 4 | 5 |
| P01034 | **1.0** | 0.9375 | **1.0** | 9 | 2 |

**Table 2:** Mean ROC-AUC performances of different variant classification models across all the amyloidosis-related proteins studied.

| Model | ROC-AUC | Completeness | Missing |
| --- | --- | --- | --- |
| AlphaMissense | $0.7612 \pm 0.1065$ | 62.5 % | P02647, P61769, P61626 |
| VESM++ | $0.7747 \pm 0.1461$ | 75% | P02647, P61769 |
| ESM2+UMAP | **$0.7851 \pm 0.1180$** | **100%** | - |

### 2.4 Classification Pipeline

The final feature matrix consisted of concatenated vectors: UMAP-reduced ESM2 embeddings. We evaluated three classifier models: logistic regression (LR), support vector machines (SVMs) with RBF kernel, and random forests. For each protein, we trained models using 5-fold crossvalidation stratified by class, and reported average precision, recall, F1-score, and area under the ROC curve (AUC).

## 3 Results

We compared our hybrid pipeline against state-of-the-art variant effect predictors such as AlphaMissense and VESM++.

**Figure 1:**
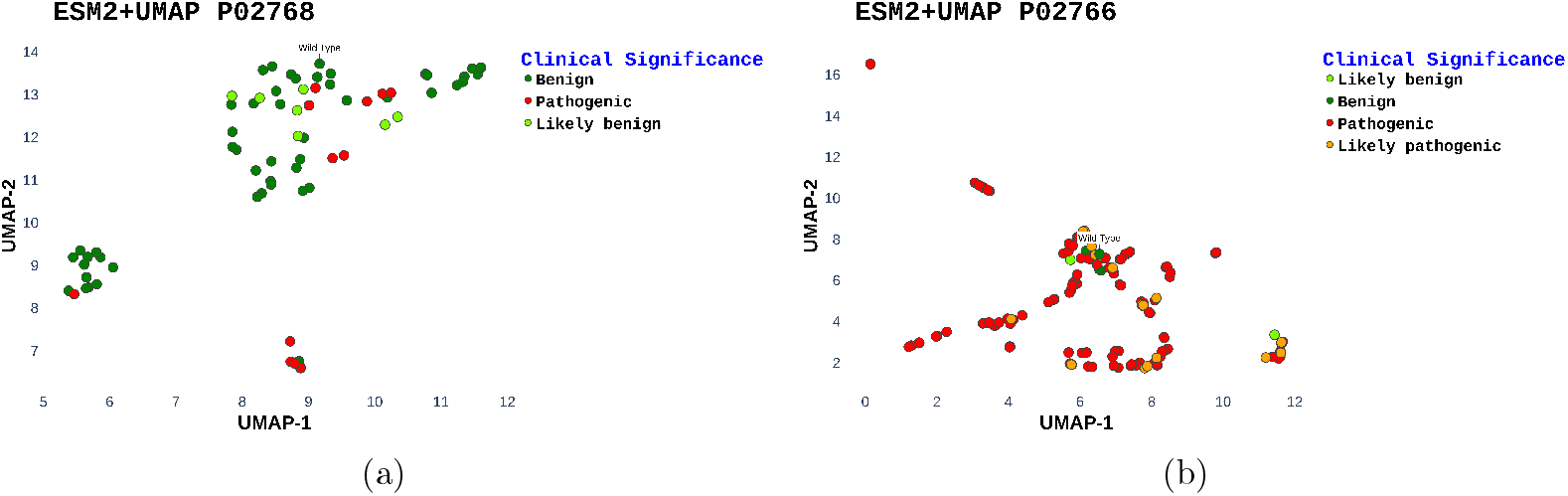
2-dimensional UMAP representation of the ESM2 variant embeddings for protein: P02768 (a) and P02766 (b).

## 4 Conclusion

This study introduces a hybrid framework that combines ESM2 protein language model (PLM) embeddings with non-linear dimensionality reduction via UMAP to support the classification and interpretation of missense variants. By leveraging the rich contextual information captured by transformer-based models and projecting these high-dimensional embeddings into a tractable space, the proposed approach enables both accurate classification and intuitive visual exploration of variant effect landscapes. Comparative analysis against state-of-the-art predictors, including AlphaMissense and VESM++, demonstrates that the ESM2+UMAP pipeline achieves competitive or superior ROC-AUC scores across a diverse set of proteins, while also offering complete coverage for all evaluated cases.

